# Exogenous butyrate inhibits butyrogenic metabolism and alters expression of virulence genes in *Clostridioides difficile*

**DOI:** 10.1101/2023.07.06.548018

**Authors:** Daniel A. Pensinger, Horia A. Dobrila, David M. Stevenson, Nicole M. Davis, Daniel Amador-Noguez, Andrew J. Hryckowian

## Abstract

The gut microbiome engenders colonization resistance against the diarrheal pathogen *Clostridioides difficile* but the molecular basis of this colonization resistance is incompletely understood. A prominent class of gut microbiome-produced metabolites important for colonization resistance against *C. difficile* is short chain fatty acids (SCFAs). In particular, one SCFA (butyrate) decreases the fitness of *C. difficile* in vitro and is correlated with *C. difficile*- inhospitable gut environments, both in mice and in humans. Here, we demonstrate that butyrate-dependent growth inhibition in *C. difficile* occurs under conditions where *C. difficile* also produces butyrate as a metabolic end product. Furthermore, we show that exogenous butyrate is internalized into *C. difficile* cells, is incorporated into intracellular CoA pools where it is metabolized in a reverse (energetically unfavorable) direction to crotonyl-CoA and (*S*)-3-hydroxybutyryl-CoA and/or 4-hydroxybutyryl-CoA. This internalization of butyrate and reverse metabolic flow of butyrogenic pathway(s) in *C. difficile* coincides with alterations in toxin production and sporulation. Together, this work highlights butyrate as a signal of a *C. difficile* inhospitable environment to which *C. difficile* responds by producing its diarrheagenic toxins and producing environmentally-resistant spores necessary for transmission between hosts. These findings provide foundational data for understanding the molecular and genetic basis of how *C. difficile* growth is inhibited by butyrate and how butyrate serves as a signal to alter *C. difficile* virulence in the face of a highly competitive and dynamic gut environment.

**IMPORTANCE:** The gut microbiome engenders colonization resistance against the diarrheal pathogen *Clostridioides difficile* but the molecular basis of this colonization resistance is incompletely understood, which hinders the development of novel therapeutic interventions for *C. difficile* infection (CDI). We investigated how *C. difficile* responds to butyrate, an end-product of gut microbiome community metabolism which inhibits *C. difficile* growth. We show that exogenously-produced butyrate is internalized into *C. difficile*, which inhibits *C. difficile* growth by interfering with its own butyrate production. This growth inhibition coincides with the expression of virulence-related genes. Future work to disentangle the molecular mechanisms underlying these growth and virulence phenotypes will likely lead to new strategies to restrict *C. difficile* growth in the gut and minimize its pathogenesis during CDI.

## INTRODUCTION

The Centers for Disease Control and Prevention classifies *Clostridioides difficile* as an “urgent threat” to the nation’s health, as it causes 450,000 infections, 15,000 deaths and 1 billion dollars in excess healthcare costs per year in the United States alone [1,2]. Dysbiosis is the primary risk factor for *C. difficile* infection (CDI) and several microbial taxa directly impact *C. difficile* fitness [3–5]. However, inter-individual variation in gut microbiome community composition complicates definitions of “CDI susceptible” versus “CDI resistant” microbiomes. Functional capacity of the distal gut microbiome (e.g., metabolites produced/consumed) differs among hosts with CDI relative to healthy hosts [6–8]. These observations from human studies and animal CDI models suggest that microbiome-dependent metabolite availability, rather than microbiome composition defines CDI susceptibility and resistance. Therefore, a focus on metabolites instead of microbes may allow for more readily translatable findings.

The gastrointestinal tract contains thousands of diverse molecules derived from the diet and host/microbiome metabolism. Many of these impact *C. difficile* fitness and pathogenesis. For example, bile acids, metals, amino acids, sugars, organic acids (OAs) and short chain fatty acids (SCFAs) affect *C. difficile in vitro* and in animal models [4–6,9–15]. SCFAs (in particular, acetate, propionate, and butyrate) are major metabolic end-products of microbiome metabolism and are primarily generated by microbial degradation of dietary fiber. Collectively, SCFAs reach concentrations >100 millimolar (mM) in bulk colon contents and are the most concentrated metabolites in the human distal gut [16]. SCFA concentrations are highly dynamic and differ with respect to microbiome composition and factors like diet, antibiotic exposure, and inflammation [17].

Much of what is known about the biology of SCFAs is based on their impacts on the host. For example, SCFAs are metabolized by colonocytes or transported systemically via portal circulation. SCFAs also enhance gut barrier function and modulate immune responses. As such, SCFA deficiencies are associated with varied diseases like inflammatory bowel disease and increased susceptibility to pathogens (Reviewed [18]). Because SCFAs are so pivotal in host-microbiome cross-talk, gut microbiome studies frequently equate a “healthy gut” with high SCFAs and a “dysbiotic gut” with low SCFAs [18]. SCFAs, in particular butyrate, inhibit *C. difficile* growth and increase *C. difficile* toxin production [6,19]. Furthermore, based on data from humans and murine models of CDI, high butyrate levels are characteristic of a gut that is non-permissive to CDI [6,15,20]. These data on CDI permissiveness, growth, and toxin production together suggest that during infection, *C. difficile* senses butyrate as a signal of a competitive environment and adjusts its virulence to maintain a dysbiosis-associated niche or to transmit to new hosts [18].

Despite the strong connections between butyrate and *C. difficile* fitness/pathogenesis and the role of this molecule as a signal of a competitive gut environment, the molecular and genetic details of how butyrate alters growth and virulence phenotypes are poorly understood. Here, using *C. difficile* 630, we show that butyrate induces large scale changes in expression of genes involved in metabolism and pathogenesis, that butyrate-dependent growth inhibition in *C. difficile* occurs under conditions where de novo butyrate synthesis pathways in *C. difficile* are active, and that exogenous butyrate is translocated into *C. difficile* cells and is metabolized through ATP- and NAD^+^-consuming pathways. Together these findings demonstrate how *C. difficile* senses and responds to butyrate and builds a strong foundation for understanding the molecular and genetic basis of these phenotypes and exploiting these processes to improve therapeutic strategies against this pathogen.

## RESULTS

### Transcriptional profiling and virulence phenotypes of butyrate-exposed *C. difficile*

To better understand butyrate-dependent growth defects in *C. difficile* [15], we performed RNA-seq on *C. difficile* 630 grown to mid-log phase (OD_600_∼0.3), early stationary phase (24 hours post inoculation), and late stationary phase (48 hours post-inoculation) in modified Reinforced Clostridial Medium (mRCM [15] supplemented with either 50 mM sodium butyrate or 50 mM sodium chloride. These media were pH adjusted to pH=6.5 to mimic the pH of the human distal gut [16] before experiments were performed (see **Methods**). The same cultures were sampled for all three time points.

We observed significant differential regulation of 313, 376, and 294 genes in response to butyrate during mid-log phase, early stationary phase, and late stationary phase, respectively (**Figure 1A, Tables S1-S3**; P<0.05, Log fold change >2). To address broader functional categories of these differentially regulated genes, we used the EGRIN models present in the *C. difficile* portal [21] to group the differentially regulated genes into functionally-related gene modules. Gene modules corresponding to metabolism, bacteriocin production, and sporulation were identified as being differentially expressed (**Figure 1A, Tables S4-S6**). While the majority of the differentially regulated genes were unique to each growth phase (**Figure 1B**), functional enrichment of metabolic genes involved in mannose and glycine fermentation were shared across all growth phases (**Table S7**). One hypothesis based on these observations and based on the metabolic plasticity of *C. difficile* [8] is that butyrate alters the metabolic preferences of *C. difficile* and that these alterations manifest as differences in growth kinetics based on nutrients available in the medium. To test this hypothesis, we supplemented mRCM with butyrate and glycine or mannose. The rationale behind these experiments is that by providing nutrients that are apparently preferred in the presence of butyrate, the growth rate of *C. difficile* would no longer be impaired by butyrate. Under these growth conditions, we did not observe a rescue of the butyrate-dependent growth defect (**Figure S1**), suggesting that other metabolic alterations may better explain butyrate-dependent growth inhibition in *C. difficile*.

**Figure 1.**
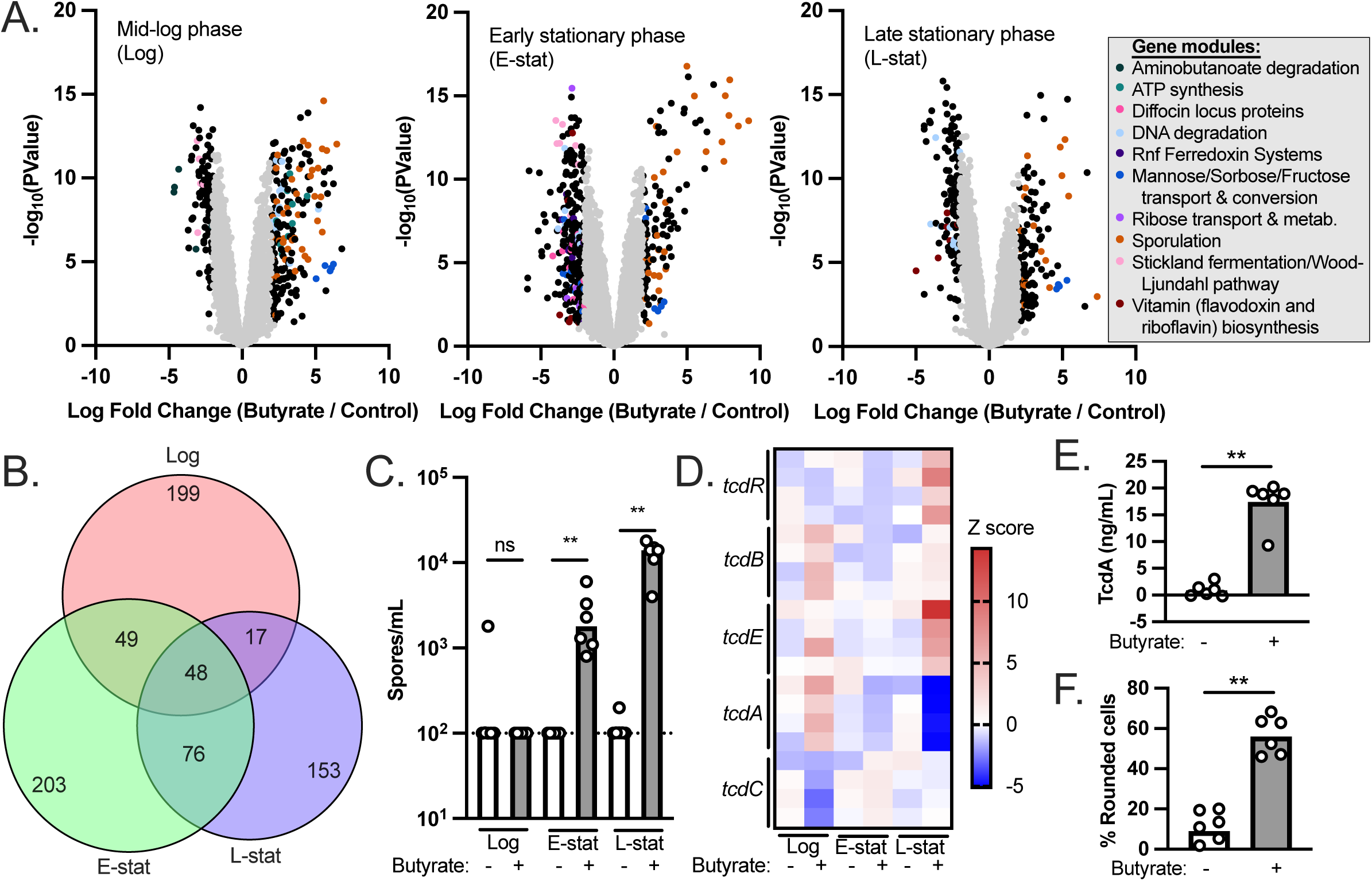
Transcriptional profiling and virulence phenotypes of butyrate-exposed *C. difficile*. (A) *C. difficile* 630 was grown to mid-log phase, early stationary phase, and late stationary phase in mRCM supplemented with either 50 mM sodium butyrate or 50 mM sodium chloride. RNA-seq was performed on n=4 independent cultures, each of which was sampled at each growth phase. Differential gene expression between sodium butyrate (butyrate) and sodium chloride (control) are shown as volcano plots for each growth phase. Functionally-related gene modules were identified among the differentially regulated genes (P<0.05, Log fold change >2) using EGRIN models present in the *C. difficile* portal [21]). (B) A Venn diagram illustrating that the majority of the significantly differentially regulated genes were unique to each growth phase. (C) Genes involved in sporulation were prominently differentially regulated genes in response to butyrate regardless of growth phase. We quantified spores in cultures of *C. difficile* during log phase, early stationary phase, and late stationary phase (n=6 independent cultures per condition, each sampled at each growth phase). (D) Previous literature demonstrates that *C. difficile* toxin production is favored in the presence of butyrate [6,19]. Transcript abundance of genes in the *C. difficile* pathogenicity locus (which contains toxin genes *tcdA* and *tcdB* as well as genes involved in toxin regulation *tcdR*, *tcdE*, and *tcdC*) were compared between sodium butyrate and sodium chloride supplemented cultures from panel A. (E) TcdA-specific ELISA demonstrates that although *tcdA* is not up-regulated at the transcriptional level, elevated levels of this toxin are present in 48-hour culture supernatants of *C. difficile* exposed to 50 mM sodium butyrate relative to 50 mM NaCl (n=6 independent cultures per condition). (F) Culture supernatants (48 hours post inoculation) of *C. difficile* 630 exposed to 50 mM sodium butyrate induce more rounding of human foreskin fibroblasts relative to culture supernatants of *C. difficile* exposed to 50 mM NaCl, indicative of increased toxin activity (n=6 independent cultures per condition). All bacterial growth media were adjusted to pH=6.5 prior to performing experiments. See also **Tables S1-S7** and **Figure S1**. **=p<0.01 via Mann-Whitney test.

In addition to the metabolic genes noted above, we observed butyrate-dependent phenotypes relating to *C. difficile* pathogenesis. First, we noted that the most abundant class of up-regulated genes, regardless of growth phase, were involved in sporulation (**Figure 1A**). To validate these gene expression data, we prepared spores from *C. difficile* cultures exposed to butyrate and observed significant butyrate-dependent increases in viable spores in butyrate-exposed cultures at early stationary phase and at late stationary phase (**Figure 1C**). Second, previous work showed that butyrate positively impacts *C. difficile* toxin production [6,19] but we did not observe a significant increase in *tcdA* or *tcdB* transcript levels at any growth phase (**Figure 1AD**). Instead, two genes within the pathogenicity locus (*tcdE* and *tcdR*) were significantly up-regulated during late stationary phase in the presence of butyrate (**Figure 1D**). To situate these counterintuitive results against previously published literature, we used two independent methods (a TcdA-specific ELISA and a cell rounding assay) to confirm that, although transcription of *tcdA* and *tcdB* are not increased in response to butyrate, the production of toxins increases in response to butyrate (**Figure 1EF**). These data demonstrate that elevated transcripts of *tcdA* and *tcdB* are not necessary for butyrate-dependent toxin production in *C. difficile* and suggest that alternative regulatory pathways may impact butyrate-dependent toxin production. While beyond the scope of the current study, an ongoing focus of our laboratory is to understand how butyrate and other host & microbiome co-metabolites influence sporulation and toxin production in *C. difficile*.

### Butyrate-dependent growth inhibition in *C. difficile* is associated with increased nutrient availability

We observed that multiple metabolic pathways are differentially regulated by *C. difficile* 630 in response to butyrate (**Figure 1AB**) but we failed to rescue butyrate-dependent growth inhibition using two high priority metabolites identified via transcriptional profiling (**Figure S1**). As exemplified by the data generated on butyrate-dependent toxin production (**Figure 1DEF**), transcriptional profiling does not capture all relevant butyrate-dependent changes to *C. difficile* gene expression. This prompted alternative approaches to investigate how exogenous butyrate impacts *C. difficile* metabolism. To begin this work, we grew *C. difficile* 630 in two base media: a complex rich medium (mRCM) and a defined minimal medium (Basal Defined Minimal Medium (BDMM) [22]). These media were supplemented with either 50 mM sodium butyrate or 50 mM sodium chloride and were pH adjusted to pH=6.5 before performing experiments as above. As expected, we observed that butyrate increases the lag time and decreases the maximum growth rate of *C. difficile* 630 in mRCM (**Figure 2A** and [15]) but we did not observe these butyrate-dependent effects on *C. difficile* 630 growth in BDMM (**Figure 2B**).

**Figure 2.**
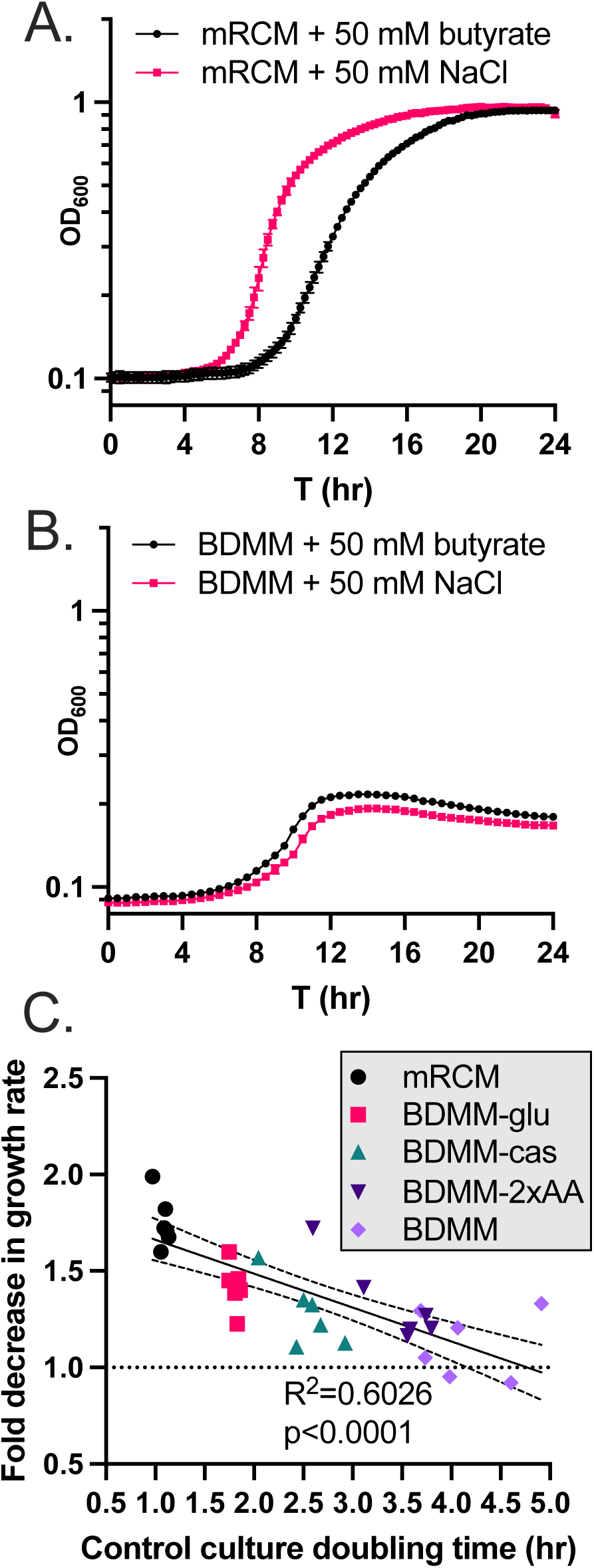
Butyrate-dependent growth inhibition in *C. difficile* as a function of growth rate. *C. difficile* 630 was grown in (A) mRCM and (B) BDMM supplemented with either 50 mM sodium butyrate or 50 mM sodium chloride (pH adjusted to pH=6.5 before performing experiments). Data points represent mean OD_600_ of 3 independent cultures (mRCM) and 5 independent cultures (BDMM) and error bars represent standard deviation. (C) The extent of growth inhibition by butyrate was determined for *C. difficile* grown in mRCM, BDMM, and three enriched BDMM formulations (BDMM-2xAA, BDMM-cas, BDMM-glu (see **Methods**)). Media were supplemented with sodium butyrate or NaCl and pH adjusted as in panels A and B. Maximum growth rates of cultures were determined using custom scripts (see **Methods**) and the growth rate of butyrate-supplemented cultures were compared against the growth rate of NaCl-supplemented cultures and a simple linear regression was carried out in GraphPad Prism, showing a significant correlation between growth rate in NaCl-supplemented cultures and degree of growth inhibition imposed by butyrate. Individual data points are shown (n=6 culture pairs per growth medium).

One hypothesis relating to these observations are that greater metabolic flux in *C. difficile*, regardless of specific nutrient sources present, is necessary for butyrate-dependent growth inhibition to occur. *C. difficile* can utilize a wide range of growth substrates and adapts its metabolism to fill a wide range of nutrient niches [23–25]. With this metabolic plasticity in mind, we addressed this hypothesis by performing growth curve analysis of *C. difficile* 630 in three enriched BDMM formulations: BDMM + 0.5% (w/v) glucose (BDMM-glu), BDMM + 1.06% (w/v) amino acids (double the concentration found in BDMM (BDMM-2x-AA)), and BDMM + 2% casamino acids in addition to the 0.53% amino acids found in BDMM (BDMM-cas). These media were supplemented with either 50 mM sodium butyrate or 50 mM sodium chloride and were pH adjusted to pH=6.5 before performing experiments as above. We found that growth of *C. difficile* 630 in these three BDMM variants was sensitive to the inhibitory effects of butyrate and that these effects significantly correlated with the increased growth rate observed in these media relative to BDMM (**Figure 2C**; R^2^=0.6026, p<0.0001). These data demonstrate that the butyrate-dependent growth defect observed in vitro in *C. difficile* occurs as a function of general nutrient availability and is not tied to the availability or abundance of any one specific nutrient source.

### *C. difficile* butyrogenic metabolism is inhibited by exogenously-applied butyrate

While exogenous butyrate inhibits the growth of diverse *C. difficile* strains under nutrient rich conditions (**Figure 2** and [15]), butyrate is a metabolic end-product of pathways in *C. difficile* that generate ATP and regenerate NAD^+^ [18]. These observations led us to hypothesize that exogenous butyrate inhibits these butyrogenic pathways in *C. difficile* via product inhibition [26]. To address this hypothesis, we quantified butyrate in supernatants from early stationary phase (24 hour) cultures of *C. difficile* 630 grown in mRCM and BDMM. In support of our hypothesis, butyrate was detected at high abundance in mRCM supernatants but was not detectable in BDMM supernatants (**Figure 3A**) and we observed a decrease in butyrate production by *C. difficile* (**Figure 3BC**) when it was grown in mRCM supplemented with butyrate. These data demonstrate that *C. difficile* produces less butyrate when in a butyrate-rich environment. Furthermore, these data suggest that that decreases in butyrogenic metabolism negatively impact its growth, perhaps by decreasing metabolic flux through key energy generating and conserving pathways.

**Figure 3.**
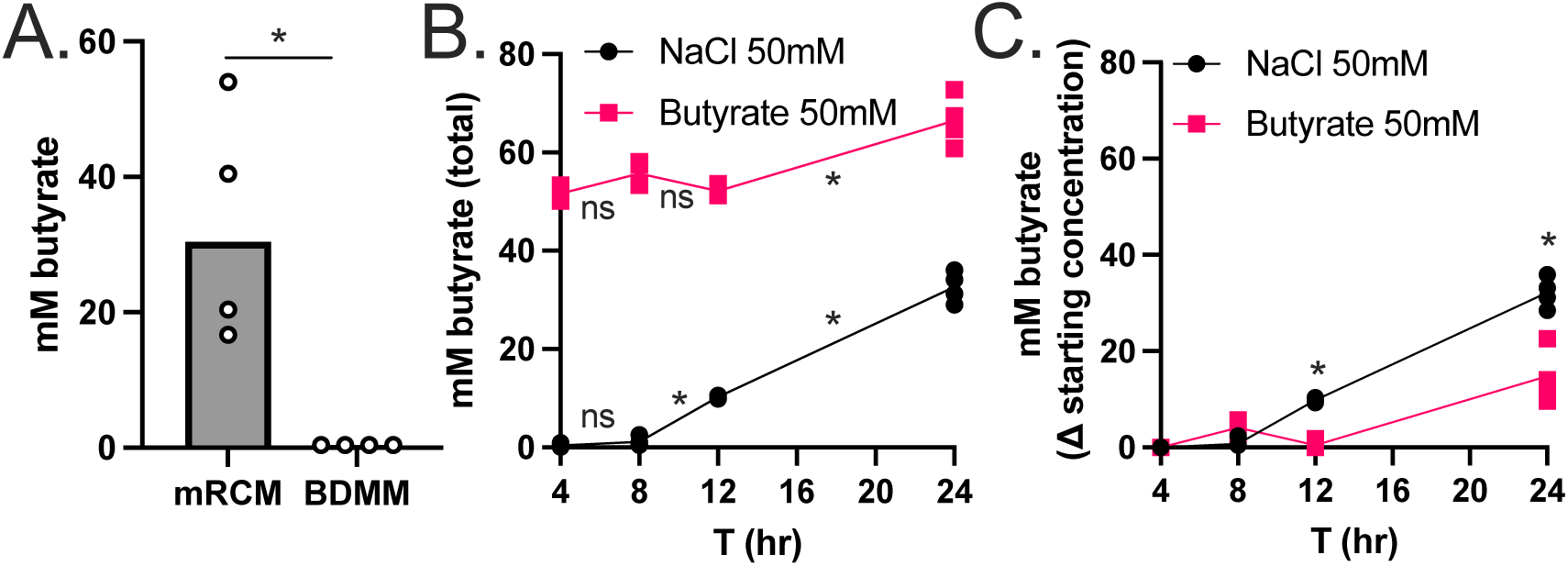
Exogenous butyrate interferes with butyrogenic metabolism in *C. difficile*. (A) Butyrate was quantified in 24-hour culture supernatants of *C. difficile* 630 grown in mRCM and BDMM via HPLC. Individual data points represent butyrate concentration and bars represent mean concentration across n=4 biological replicates per condition. (B) Total butyrate was quantified in culture supernatants of *C. difficile* 630 grown in mRCM supplemented with 50 mM sodium butyrate or 50 mM NaCl at 4, 8, 12, and 24 hours post inoculation. (C) *C. difficile* butyrate production was determined for butyrate-supplemented cultures by subtracting 50 mM from the total amount of butyrate detected in panel B. All media were adjusted to pH=6.5 prior to use in experiments. For panels B and C, individual data points represent mean concentration and lines connect mean concentrations over time across n=4 biological replicates per condition. Statistical significance was determined by Mann-Whitney test. *=p<0.05

### Extracellular butyrate is internalized into *C. difficile*, is incorporated into CoA pools, and is metabolized in an energetically unfavorable direction

To better understand the impacts of butyrate on *C. difficile* metabolism, we quantified intracellular pools of butyryl-CoA in *C. difficile* 630 grown in 25 mM sodium butyrate relative to *C. difficile* 630 grown in 25 mM NaCl during log-phase growth via LC/MS. We observed elevated levels of intracellular butyrate, in the form of butyryl-CoA, in butyrate-supplemented cultures relative to NaCl-supplemented cultures (**Figure 4A**). These data suggest either that *C. difficile*-produced butyrate is not secreted into the extracellular environment or that exogenous butyrate enters *C. difficile* cells and is incorporated into intracellular CoA pools.

**Figure 4.**
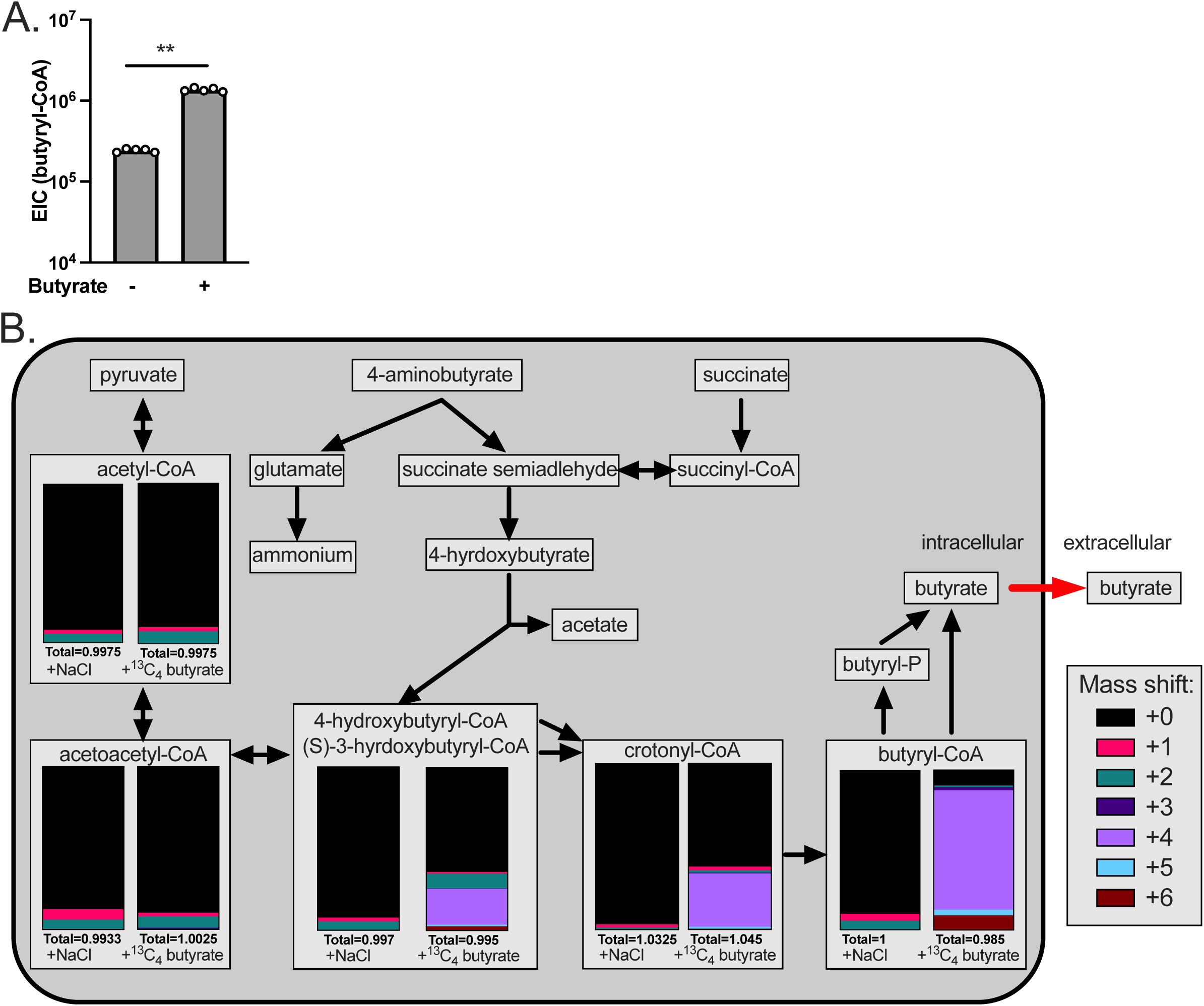
Exogenous butyrate is internalized into *C. difficile* cells, is incorporated into CoA pools, and forces energetically unfavorable metabolic processes. (A) *C. difficile* was grown in mRCM supplemented with 25 mM sodium butyrate or 25 mM NaCl and intracellular butyryl-CoA levels were quantified by LC/MS from cells collected at mid-log phase. Individual data points represent abundance of butyryl-CoA and bars represent mean abundance across n=5 biological replicates per condition. (B) *C. difficile* was grown in mRCM supplemented with 25 mM ^13^C_4_ sodium butyrate or 25 mM NaCl and isotopically labeled CoA intermediates were quantified at mid-log phase. CoA intermediates detectable by the LC/MS method under the ^13^C_4_ sodium butyrate- and NaCl- supplemented conditions are overlaid onto a metabolic map of butyrogenic pathways in *C. difficile* inferred from BioCyc [47] and arrows indicate energetically-favorable directionality of the reactions. Stacked bar charts represent the mean abundance of molecules with corresponding mass shifts as annotated in the figure key (n=4 biological replicates per condition). All media were adjusted to pH=6.5 prior to use in experiments. Statistical significance was determined by Mann-Whitney test. **=p<0.01.

To determine whether the increase in intracellular butyryl-CoA in *C. difficile* was due to elevated levels of endogenously produced butyrate that fails to be released or due to import of extracellular butyrate, we supplemented mRCM with 25mM ^13^C_4_ sodium butyrate and examined metabolic incorporation of the ^13^C label into *C. difficile* CoA pools by LC-MS. In addition to ^13^C_4_- butyryl-CoA, we also observed increased relative abundance of ^13^C_4_-crotonyl-CoA in ^13^C_4_ sodium butyrate-exposed C. difficile relative to 25 mM NaCl-supplemented controls (**Figure 4B).** In addition, we observed significant enrichment of molecule(s) with m/z=853.152 containing the ^13^C_4_ label. Molecules at this m/z correspond to (*S*)-3-hydroxybutyryl-CoA and/or 4-hydroxybutyryl-CoA (**Figure 4B**), which are CoA intermediates upstream of crotonyl-CoA in the butyrogenic pathways present in *C. difficile* (**Figure 4B**). Importantly, (S)-3-hydroxybutyryl-CoA and/or 4-hydroxybutyryl-CoA are isomers and cannot be resolved with LC/MS and other CoA intermediates further upstream in these pathways were either present below the limit of detection (e.g., succinyl-CoA) or did not have ^13^C label above background (e.g., acetoacetyl-CoA & acetyl-CoA) (**Figure 4B**). Regardless, the enrichment of ^13^C label in butyryl-CoA, crotonyl-CoA, and (*S*)-3-hydroxybutyryl-CoA/4-hydroxybutyryl-CoA indicates that exogenous butyrate is internalized into *C. difficile* cells and drives butyrogenic reactions in reverse (energetically unfavorable) directions in *C. difficile,* likely contributing to its reduced fitness in a butyrate-rich environment.

## DISCUSSION

This work provides several insights into how *C. difficile* responds to butyrate, a prominent gut microbiome produced metabolite. Specifically, we demonstrate broad changes to the expression of metabolic and virulence genes in response to butyrate (**Figure 1**), that the growth inhibition observed in *C. difficile* in response to butyrate occurs under nutrient rich conditions where *C. difficile* produces butyrate from its own butyrogenic pathways (**Figures 2 and 3**). Furthermore, we show that exogenous butyrate is internalized into *C. difficile* cells, is incorporated into intracellular CoA pools where it is metabolized in a reverse (energetically unfavorable) direction to crotonyl-CoA and (*S*)-3-hydroxybutyryl-CoA and/or 4-hydroxybutyryl-CoA (**Figure 4**). These data support a model where *C. difficile* butyrogenic metabolism is negatively impacted by butyrate internalized from the environment and that these effects collectively lead to the butyrate-dependent growth defects and butyrate-dependent colonization resistance observed in this and previous work [6,15]. Furthermore, this work provides possible explanations for inconsistencies in previous literature, where butyrate rich environments are not always associated with reduced *C. difficile* fitness [27,28]. Specifically, because butyrate-dependent growth inhibition occurs under conditions where *C. difficile* produces butyrate (**Figure 3A**), it is possible that these previous studies did not provide the necessary environment (e.g., growth substrates and butyrate) for fitness to be negatively impacted. Continued investigation of how *C. difficile* responds to butyrate will shed light on the conditions that impact its pleiotropic effects on *C. difficile* and can serve as a model of how specific molecules, against the backdrop of a variable & complex metabolic milieu, may have differential impacts on gut resident microbes.

Beyond metabolism, the response of *C. difficile* to a nutrient- and butyrate-rich metabolic environment provides key insights into butyrate as a signaling molecule for *C. difficile* and highlights paths for future research. First, we observed increased toxin production by *C. difficile* in response to butyrate, as in previous literature that used both murine models of infection and *in vitro* approaches (**Figure 1EF** and [6,19]). Although a complex regulatory network controls the production of *C. difficile* toxins [29], our work highlights additional uncharacterized levels of control of toxin production in *C. difficile*. Specifically, we did not observe increases in *tcdA* or *tcdB* transcripts under any growth phase (**Figure 1A, 1D, Tables S1-S3**) but did observe increased levels of TcdA by ELISA and increased biologically active toxin (a combination of TcdA and TcdB) by a cell rounding assay (**Figure 1EF**). These combined observations suggest butyrate-dependent post-transcriptional regulation of *C. difficile* toxins. Second, we observed butyrate-dependent increases in sporulation-related genes at all growth phases and in viable spores during stationary phase (**Figure 1A-C**). While numerous regulators of sporulation in *C. difficile* have been characterized [30–33], the specific effects of butyrate on sporulation remain uncharacterized. While beyond the scope of the current study, an ongoing focus of our laboratory is to understand how butyrate and other host & microbiome co-metabolites influence sporulation and toxin production in *C. difficile*.

The model that *C. difficile* senses butyrate as a signal of a competitive and inhospitable gut environment fits with long established paradigms of diverse pathogenic bacteria enacting virulence and transmission programs under unfavorable environmental conditions (e.g., *C. difficile*, *Enterococcus*, and *Salmonella* [34–36]. In the context of our *in vitro* observations of *C. difficile* described here, we expect that the decreased fitness and increased toxin production and sporulation to be relevant under conditions when *C. difficile* is abundant in the gut and begins to face increasingly abundant competing microbes as the microbiome recovers from dysbiosis, such as after cessation of antibiotic treatment [37] or after a change in diet that results in abundant butyrate production by the gut microbiome [15]. This may simultaneously allow *C. difficile* to cause inflammation and compete with inflammation-sensitive members of the microbiome, as has been observed for pathogens like *Salmonella* [38] and facilitate transmission of spores to new hosts. In the context of a healthy microbiome where *C. difficile* is unable to colonize, we expect that the effects of growth inhibition would dominate and contribute to colonization resistance against the pathogen and facilitate spore formation.

While this work focuses on *C. difficile*, it is unlikely that similar butyrate-dependent effects are seen all Clostridia – both due to differences in gene content and ecological niche. A recent example illustrating this idea explored butyrate sensitivity in *Bacteroides* strains and showed that *Bacteroides thetaiotaomicron* could be engineered to be susceptible to butyrate by altering the acyl-CoA thioesterase to be less efficient at disassociating butyrate from butyryl-CoA [39]. In this previous work, the acyl-CoA thioesterase from *B. thetaiotaomicron* (butyrate-resistant) was replaced with the acyl-CoA thioesterase from butyrate-sensitive *Bacteroides vulgatis.* Both butyrate sensitivity and a buildup of butyryl-CoA were associated with this mutant strain. The authors of that study hypothesized that such a “backup” of butyryl-CoA had an effect of sequestering the CoA molecule. In the context of *C. difficile*, it is possible that its acyl-CoA thioesterase(s) are inefficient and this manifests as an increase in butyryl-CoA and energetically unfavorable flux through up-stream CoA intermediates. While beyond the scope of our study, it remains to be determined if other organisms related to *C. difficile* have alleles of acyl-CoA thioesterase (or other gene(s)) that render them insensitive to the growth inhibitory effects of butyrate. Furthermore, finer-scale dissection of which specific aspect of the butyrate pathway disruption (energetically unfavorable metabolism of butyrate, lack of NADH turnover, backup of CoA pools, or other effects) are the major factor involved in butyrate-dependent growth inhibition and which of these effects serves as a signal to *C. difficile* to alter the expression of genes involved in pathogenesis.

In summary, our work further establishes butyrate as an important signal to *C. difficile* of the “health” of the microbiome by directing its metabolic and virulence programs. This view fits with an increasing appreciation of metabolites as signaling molecules influencing physiological processes outside of metabolism [40]. In the case of *C. difficile,* exogenous butyrate affects the functioning of its own butyrate synthesis pathways and elicits a response from the bacterium to increase the inflammatory state of the gut and produce environmentally-resistant spores necessary for transmission.

## METHODS

### Bacterial strains and culture conditions

All bacterial growth media were pre-reduced for a minimum of 24 hours in an anaerobic chamber prior to use in experiments and all bacterial growth was done under anaerobic conditions in an anaerobic chamber (Coy).

*C. difficile* 630 was maintained as −80°C stocks in 25% glycerol under anaerobic conditions in septum-topped vials. *C. difficile* was routinely cultured on CDMN agar (*Clostridioides difficile* agar with moxalactam and norfloxacin), composed of *C. difficile* agar base (Oxoid) supplemented with 7% defibrinated horse blood (HemoStat Laboratories), 32 mg/L moxalactam (Santa Cruz Biotechnology), and 12 mg/L norfloxacin (Sigma-Aldrich) in an anaerobic chamber at 37°C (Coy). After 16 to 24 h of growth on agar plates under anaerobic conditions, a single colony was picked into 5 mL of modified reinforced clostridial medium (mRCM; [15] or BDMM [11] and grown anaerobically at 37°C for 16 to 24 h.

For *in vitro* growth curve experiments examining bacterial growth, subcultures were prepared at 1:200 dilutions in mRCM and BDMM supplemented with sodium butyrate, sodium chloride, glycine, mannose, glucose, amino acids, and casamino acids as specified in the main text and in the figure legends. After addition of these media supplements, the pH of the media was adjusted as specified in the figure legends. Growth curve experiments were done in sterile polystyrene 96-well tissue culture plates with low evaporation lids (Falcon). Cultures were grown anaerobically in a BioTek Epoch2 plate reader. At 15- or 30-minute intervals, the plate was shaken on the “slow” setting for 1 min and the OD_600_ of the cultures was recorded using Gen5 software (version 1.11.5).

### Transcriptional profiling of *C. difficile* in response to butyrate

*C. difficile* 630 overnight cultures were back-diluted 1:50 in 20mL mRCM in 50mL conical tubes with 50 mM sodium butyrate or 50 mM sodium chloride adjusted to pH=6.5. Harvest was performed at OD_600_=0.3-0.5 for mid-log phase samples, at 24 hours post inoculation (early stationary phase) or at 48 hours post inoculation (late stationary phase).

At the appropriate time points, 5mL aliquots of the cultures were diluted 1:1 in chilled 1:1 Ethanol:Acetone to preserve RNA [41]. These samples were then centrifuged for 5 minutes at 3000 x g at room temperature and cell pellets were frozen at −80°C. Immediately prior to RNA extraction, cell pellets were centrifuged at 4°C at 3000 x g for 1 minute. Residual supernatant was removed from the cell pellets, which were subsequently washed with 5mL nuclease free PBS. Washed pellets were centrifuged at 4°C at 3000 x g for 1 minute, the supernatant was removed, and the resulting pellet was resuspended in 1mL TRIzol and processed using a TRIzol Plus RNA Purification Kit (Thermo) with on-column DNase treatment according to the manufacturer’s instructions. Purified RNA was frozen at −80C and RNA integrity was confirmed via BioAnalyzer (Agilent) prior to proceeding.

RNA-seq on high quality rRNA-depleted RNA extracts (12M paired end reads per sample) and transcript level quantification, count normalization, and differential expression analysis were performed using the *C. difficile* 630 reference genome (GCF_000009205.2_ASM920v2) for sequence alignments (SeqCenter, Pittsburgh, Pennsylvania, USA). See also **Data Availability**.

### *C. difficile* spore quantification

Overnight cultures of *C. difficile* were diluted 1:200 into pre-reduced mRCM supplemented with either 50mM sodium butyrate or 50mM NaCl (pH adjusted to pH=6.5) and incubated anaerobically at 37°C. At 8, 24, and 48 hours post inoculation, 0.5 mL aliquots of each culture were taken, heated to 65°C for 20 mins to kill vegetative cells, and 10-fold serially diluted in pre-reduced mRCM. Ten microliters of the serial dilutions were then spotted onto pre-reduced RCM plates containing 0.1% sodium taurocholate. Dilutions were also plated onto pre-reduced RCM plates without 0.1% sodium taurocholate to confirm no vegetative cells were present. Plates were incubated for 48 hours at 37°C and colonies present on RCM plates containing 0.1% sodium taurocholate were quantified. No colonies from heat-treated samples were observed on RCM plates without taurocholate.

### *C. difficile* toxin quantification

*C. difficile* toxin was quantified in 48-hour culture supernatants using two assays. First, levels of TcdA were quantified relative to a standard curve of purified TcdA using the Separate Detection of *C. difficile* Toxins A and B Kit (TGCBiomics) according to the manufacturer’s instructions.

Second, a cell rounding assay was performed (modified from [42]). Two days before cell treatment, overnight cultures of *C. difficile* grown in mRCM were back-diluted 1:200 into mRCM containing 50 mM sodium butyrate or 50 mM sodium chloride (pH adjusted to pH=6.5). Cultures were grown anaerobically at 37°C for 48 hours with 50mM butyrate or NaCl. One day before treatment, confluent human foreskin fibroblast (HFF) cells (ATCC SCRC-1041) grown in HFF medium (see below for HFF medium composition) were harvested and counted with a hemocytometer and seeded into 48-well plates at 15,000 cells per well and incubated at 37°C under 5% CO_2_ for 24 hours. On the day of experiments, *C. difficile* culture supernatants were spun (3,000 x g for 15 minutes at room temperature) and were filter sterilized.

Because butyrate reduces cytotoxicity of *C. difficile* toxins on eukaryotic cells [27], we removed butyrate from culture supernatants by size exclusion filtration to minimize these effects and our interpretation of the data. Specifically, for each culture, 5 mL of filtered culture supernatant was transferred to Amicon 100 kDa MWCO filters and was centrifuged at 3,000 x g for 5 mins at room temperature. The filter dead volume (containing molecules >100 kDa, including *C. difficile* toxins) was then washed with 5 mL of 1X PBS and was centrifuged again, as above. Then, the washed fraction containing *C. difficile* toxins was reconstituted to 5 mL with HFF medium, in order to dilute this fraction back to original concentration found in culture supernatants. Concentrations of the resulting toxin-containing HFF media were prepared at: undiluted, 1:50, 1:100, and 1:150 dilutions. Then, phase contrast images were acquired using Sartorius Incucyte to confirm the health and morphology of the cells prior to treatment. Then, the medium was removed from the 24 hour-grown HFF cells and toxin-containing media at the dilutions noted above were added. Cells were incubated with toxin for 4 hours and imaged again. Total and rounded cells were counted manually. While all dilutions of toxin-containing media showed significant differences between butyrate- and NaCl-supplemented conditions (data not shown), the 1:100 dilution of toxin-containing HFF medium was used to generate the data in **Figure 1F**.

HFF medium contains the following components by volume: Eighty-eight percent high glucose DMEM (Thermo Scientific 11965126), 10% heat-inactivated fetal bovine serum, 1% penicillin/streptomycin (Thermo Scientific 15140122), 1% Glutamax (L-alanyl-L-glutamine) (Thermo Scientific 35050061).

### Measurement of maximum growth rate for in vitro growth experiments

Raw OD_600_ measurements of bacterial cultures (see **Bacterial strains and culture conditions**) were exported from Gen5 and analyzed as previously described [15]. Growth rates were determined for each culture by calculating the derivative of natural log-transformed OD_600_ measurements over time. Growth rate values at each time point were then smoothed using a moving average over 90-min intervals to minimize artifacts due to noise in OD measurement data, and these smooth growth rate values were used to determine the maximum growth rate for each culture. To mitigate any remaining issues with noise in growth rate values, all growth rate curves were also inspected manually. Specifically, in cases where the growth_curve_statistics.py script selected an artifactual maximum growth rate, the largest local maximum that did not correspond to noise was manually assigned as the maximum growth rate.

### HPLC-based quantification of butyrate in culture supernatants

Butyrate was quantified in bacterial culture supernatants as previously described [43]. Overnight cultures of *C. difficile* 630 grown in mRCM and sub-cultured into mRCM with 50 mM of butyrate and NaCl bacterial cultures. At the time points specified in the figures, cultures were centrifuged at 3000 x g for 5 minutes and the resulting supernatant was collected, 0.22µm filtered, and stored −20 °C. Supernatants were thawed and H_2_SO_4_ was added to a final concentration of 18 mM. Samples were mixed, incubated 2 minutes at room temperature and centrifuged at 21,000 x g for 10 minutes at 4°C. Soluble fractions were aliquoted into HPLC vials. In addition, 100 mM, 20 mM, and 4 mM butyrate standards were prepared in mRCM or BDMM, as appliable and processed as above. HPLC analysis was performed with a ThermoFisher (Waltham, MA) Ultimate 3000 UHPLC system equipped with a UV detector (210 nm). Compounds were separated on a 250 × 4.6 mm Rezex© ROA-Organic acid LC column (Phenomenex Torrance, CA) run with a flow rate of 0.3 mL min−1 and at a column temperature of 50 °C. Separation was isocratic with a mobile phase of HPLC grade water acidified with 0.015 N H_2_SO_4_. Resulting data were analyzed with ThermoFisher Chromeleon 7 and butyrate concentrations in culture supernatants were determined by analysis against a standard curve as described above.

### LC/MS-based CoA-targeted intracellular metabolomics

Overnight cultures of *C. difficile* grown in mRCM were back-diluted 1:50 in 20mL mRCM supplemented with 25 mM sodium butyrate, 25 mM ^13^C_4_ sodium butyrate, or 25 mM NaCl in 50mL conical tubes and incubated 37°C. The pH of mRCM used in these experiments was at the natural pH of the media (typically 6.5-7) for the experiments using sodium butyrate and the pH of the media for the^13^C_4_ sodium butyrate experiments was adjusted to pH=6.5 as needed. Cultures harvested at mid-log phase anaerobically −5mL of culture was deposited by vacuum filtration onto a 0.2 µm nylon membrane (47 mm diameter) in duplicate. The membrane was then placed (cells down) into 1.5 ml cold (on dry ice) extraction solvent (20:20:10 v/v/v acetonitrile, methanol, water) in a small petri dish and swirled. After approximately 30 seconds, the filter was inverted (cells up) and solvent was passed over the surface of the membrane several times to maximize extraction. The cell extract was then stored at −80°C. Prior to LC/MS analysis, extracts were centrifuged at 21,000 x g at 4°C for 10 minutes. Next, ∼200μL of extract normalized to OD_600_ was dried over N_2_ gas. Extracts were resuspended in 70μL of HPLC grade water and pelleted at 21,000 x g at 4°C for 10 minutes to remove particulates. All cultures were extracted in biological triplicate or quadruplicate.

For experiments using non-labeled sodium butyrate, extracts were analyzed by mass spectrometry as previously described except without MOPS exclusion [44]. Briefly, samples were run through an ACQUITY UPLC® BEH C18 column in an 18-minute gradient with Solvent A being 97% water, 3% methanol, 10 mM tributylamine (TBA), 9.8 mM acetic acid, pH 8.2 and Solvent B being 100% methanol. The gradient was 5% Solvent B for 2.5 minutes, gradually increased to 95% Solvent B at 18 minutes, held at 95% Solvent B until 20.5 minutes, returned to 5% Solvent B over 0.5 minutes, and held at 5% Solvent B for the remaining 4 minutes. Ions were generated by heated electrospray ionization (HESI; negative mode) and quantified by a hybrid quadrupole high-resolution mass spectrometer (Q Exactive orbitrap, Thermo Scientific). MS scans consisted of full MS scanning for 70-1000 *m/z* from time 0–18 min. Metabolite peaks were identified using Metabolomics Analysis and Visualization Engine (MAVEN) [45,46].

For experiments using ^13^C_4_ sodium butyrate, the protocol was adjusted to increase resolution of CoA conjugated molecules with the following modifications. The m/z window was adjusted to exclude <300 Daltons, the first 5 minutes of run was excluded, and injection volume was increased from 10µL to 25µL.

### Data availability

Data on normalized transcript abundance and differential expression analysis are found in **Tables S1-S3**. Prior to publication of a peer-reviewed manuscript, the raw data from the RNA-seq experiments shown in **Figure 1** and **Tables S1-S3** will be available from the corresponding author upon request. These raw data will be uploaded to NCBI and made freely available upon acceptance of the peer-reviewed manuscript.

### Statistical analysis

Statistical analysis was performed using GraphPad Prism 9.1.0. Details of specific analyses, including statistical tests used, are found in applicable figure legends. * = p<0.05, ** = p<0.01, *** = p<0.005, **** = p<0.001.

## Acknowledgments

We thank Cody Martin for technical assistance with bacterial growth rate calculations. Research reported in this publication was supported by startup funding from the University of Wisconsin-Madison (AJH), and a gift from Judy and Sal Troia (AJH), and an NIH NRSA award (F32AI172084 to NMD).

## Author contributions

DAP, HAD, and NMD performed experiments and analyzed the data. DAP, HAD, and AJH prepared the display items. DMS, NMD, DAN, and AJH provided key insights, tools, and reagents. DAP and AJH wrote the paper. All authors edited and approved the manuscript prior to submission.

## Figure Legends

<u>Figure S1. *C. difficile* butyrate-dependent growth defects are not rescued in mRCM</u> <u>supplemented with mannose or glycine. (A)</u> *C. difficile* 630 was grown in mRCM supplemented with 100 mM sodium butyrate or with 100 mM sodium butyrate + 50 mM glycine. (B) *C. difficile* 630 was grown in mRCM supplemented with 50 mM sodium butyrate or with 50 mM sodium butyrate + 50 mM mannose. All media were adjusted to pH=6.5 prior to use in experiments. Data points represent mean OD_600_ of three independent cultures and error bars represent standard deviation. Related to Figure 1.

<u>Table S1. Genes differentially regulated by *C. difficile* 630 in response to butyrate during mid-log</u> <u>phase.</u> Related to **Figure 1**.

<u>Table S2. Genes differentially regulated by *C. difficile* 630 in response to butyrate during early</u> <u>stationary phase.</u> Related to **Figure 1**.

<u>Table S3. Genes differentially regulated by *C. difficile* 630 in response to butyrate during late</u> <u>stationary phase.</u> Related to **Figure 1**.

<u>Table S4. Gene modules enriched within the genes differentially regulated by *C. difficile* 630 in</u> <u>response to butyrate during mid-log phase.</u> Gene modules were defined using the *C. difficile* portal [21]. Related to **Figure 1** and **Table S1.**

<u>Table S5. Gene modules enriched within the genes differentially regulated by *C. difficile* 630 in</u> <u>response to butyrate during early stationary phase.</u> Gene modules were defined using the *C. difficile* portal [21]. Related to **Figure 1** and **Table S2.**

<u>Table S6. Gene modules enriched within the genes differentially regulated by *C. difficile* 630 in</u> <u>response to butyrate during late stationary phase.</u> Gene modules were defined using the *C. difficile* portal [21]. Related to **Figure 1** and **Table S3.**

<u>Table S7. Gene modules enriched within genes differentially regulated by *C. difficile* 630 in</u> <u>response to butyrate across all growth phases.</u> Gene modules were defined for genes that were differentially regulated across all growth phases using the *C. difficile* portal [21]. Related to **Figure 1** and **Tables S1-S3.**

## REFERENCES

1. Guh AY, Mu Y, Winston LG, Johnston H, Olson D, Farley MM, et al. Trends in U.S. Burden of *Clostridioides difficile* Infection and Outcomes. N Engl J Med. N Engl J Med; 2020;382: 1320–1330. doi:10.1056/NEJMOA1910215 PMID:32242357

2. Lessa FC, Mu Y, Bamberg WM, Beldavs ZG, Dumyati GK, Dunn JR, et al. Burden of *Clostridium difficile* Infection in the United States . N Engl J Med. 2015;372: 825–834. doi:10.1056/nejmoa1408913 PMID:25714160

3. Schubert AM, Sinani H, Schloss PD. Antibiotic-Induced Alterations of the Murine Gut Microbiota and Subsequent Effects on Colonization Resistance against *Clostridium difficile*. MBio. mBio; 2015;6. doi:10.1128/MBIO.00974-15 PMID:26173701

4. Ferreyra JA, Wu KJ, Hryckowian AJ, Bouley DM, Weimer BC, Sonnenburg JL. Gut microbiota-produced succinate promotes *C. difficile* infection after antibiotic treatment or motility disturbance. Cell Host Microbe. Cell Host Microbe; 2014;16: 770–777. doi:10.1016/J.CHOM.2014.11.003 PMID:25498344

5. Buffie CG, Bucci V, Stein RR, McKenney PT, Ling L, Gobourne A, et al. Precision microbiome reconstitution restores bile acid mediated resistance to *Clostridium difficile*. Nature. Nature; 2015;517: 205–208. doi:10.1038/NATURE13828 PMID:25337874

6. Hryckowian AJ, Van Treuren W, Smits SA, Davis NM, Gardner JO, Bouley DM, et al. Microbiota-accessible carbohydrates suppress *Clostridium difficile* infection in a murine model. Nat Microbiol. Nat Microbiol; 2018;3: 662–669. doi:10.1038/S41564-018-0150-6 PMID:29686297

7. Rojo D, Gosalbes MJ, Ferrari R, Pérez-Cobas AE, Hernández E, Oltra R, et al. *Clostridium difficile* heterogeneously impacts intestinal community architecture but drives stable metabolome responses. ISME J. ISME J; 2015;9: 2206–2220. doi:10.1038/ISMEJ.2015.32 PMID:25756679

8. Jenior ML, Leslie JL, Young VB, Schloss PD. *Clostridium difficile* Colonizes Alternative Nutrient Niches during Infection across Distinct Murine Gut Microbiomes. mSystems. American Society for Microbiology; 2017;2. doi:10.1128/msystems.00063-17

9. Zackular JP, Moore JL, Jordan AT, Juttukonda LJ, Noto MJ, Nicholson MR, et al. Dietary zinc alters the microbiota and decreases resistance to *Clostridium difficile* infection. Nat Med. Nat Med; 2016;22: 1330–1334. doi:10.1038/NM.4174 PMID:27668938

10. Fletcher JR, Pike CM, Parsons RJ, Rivera AJ, Foley MH, McLaren MR, et al. *Clostridioides difficile* exploits toxin-mediated inflammation to alter the host nutritional landscape and exclude competitors from the gut microbiota. Nat Commun. Nat Commun; 2021;12. doi:10.1038/S41467-020-20746-4 PMID:33469019

11. Battaglioli EJ, Hale VL, Chen J, Jeraldo P, Ruiz-Mojica C, Schmidt BA, et al. *Clostridioides difficile* uses amino acids associated with gut microbial dysbiosis in a subset of patients with diarrhea. Sci Transl Med. Sci Transl Med; 2018;10. doi:10.1126/SCITRANSLMED.AAM7019 PMID:30355801

12. Collins J, Robinson C, Danhof H, Knetsch CW, Van Leeuwen HC, Lawley TD, et al. Dietary trehalose enhances virulence of epidemic *Clostridium difficile*. Nature. Nature; 2018;553: 291–294. doi:10.1038/NATURE25178 PMID:29310122

13. Pruss KM, Sonnenburg JL. *C. difficile* exploits a host metabolite produced during toxin-mediated disease. Nature. Nature; 2021;593: 261–265. doi:10.1038/s41586-021-03502-6

14. Pruss KM, Enam F, Battaglioli E, DeFeo M, Diaz OR, Higginbottom SK, et al. Oxidative ornithine metabolism supports non-inflammatory *C. difficile* colonization. Nat Metab. Nat Metab; 2022;4: 19–28. doi:10.1038/S42255-021-00506-4 PMID:34992297

15. Pensinger DA, Fisher AT, Dobrila HA, Treuren W Van, Gardner JO, Higginbottom SK, et al. Butyrate Differentiates Permissiveness to *Clostridioides difficile* Infection and Influences Growth of Diverse *C. difficile* Isolates. Infect Immun. 2023;91: 1–13. doi:10.1128/iai.00570-22 PMID:36692308

16. Cummings JH, Pomare EW, Branch HWJ, Naylor CPE, MacFarlane GT. Short chain fatty acids in human large intestine, portal, hepatic and venous blood. Gut. Gut; 1987;28: 1221–1227. doi:10.1136/GUT.28.10.1221 PMID:3678950

17. Macfarlane S, Macfarlane GT. Regulation of short-chain fatty acid production. Proc Nutr Soc. Proc Nutr Soc; 2003;62: 67–72. doi:10.1079/PNS2002207 PMID:12740060

18. Gregory AL, Pensinger DA, Hryckowian AJ. A short chain fatty acid-centric view of *Clostridioides difficile* pathogenesis. PLoS Pathog. PLoS Pathog; 2021;17. doi:10.1371/JOURNAL.PPAT.1009959 PMID:34673840

19. Karlsson S, Lindberg A, Norin E, Burman LG, Akerlund T. Toxins, butyric acid, and other short-chain fatty acids are coordinately expressed and down-regulated by cysteine in *Clostridium difficile*. Infect Immun. American Society for Microbiology; 2000;68: 5881– 5888. PMID:10992498

20. Seekatz AM, Theriot CM, Rao K, Chang YM, Freeman AE, Kao JY, et al. Restoration of short chain fatty acid and bile acid metabolism following fecal microbiota transplantation in patients with recurrent *Clostridium difficile* infection. Anaerobe; 2018;53: 64–73. doi:10.1016/J.ANAEROBE.2018.04.001 PMID:29654837

21. Arrieta-Ortiz ML, Immanuel SRC, Turkarslan S, Wu W-J, et al. Predictive regulatory and metabolic network models for systems analysis of *Clostridioides difficile*. Cell Host Microbe. 2021;29: 1709–1723. doi:10.1016/j.chom.2021.09.008

22. Karasawa T, Ikoma S, Yamakawa K, Nakamura S. A defined growth medium for *Clostridium difficile*. Microbiology. Microbiology (Reading); 1995;141 (Pt 2): 371–375. doi:10.1099/13500872-141-2-371 PMID:7704267

23. Neumann-Schaal M, Jahn D, Schmidt-Hohagen K. Metabolism the difficile way: The key to the success of the pathogen *Clostridioides difficile*. Frontiers in Microbiology. Frontiers Media S.A.; 2019. p. 219. doi:10.3389/fmicb.2019.00219

24. Jenior ML, Leslie JL, Young VB, Schloss PD. *Clostridium difficile* Colonizes Alternative Nutrient Niches during Infection across Distinct Murine Gut Microbiomes . mSystems. American Society for Microbiology; 2017;2.

25. Riedel T, Wetzel D, Hofmann JD, Plorin SPEO, Dannheim H, Berges M, et al. High metabolic versatility of different toxigenic and non-toxigenic *Clostridioides difficile* isolates. Int J Med Microbiol. Int J Med Microbiol; 2017;307: 311–320. doi:10.1016/J.IJMM.2017.05.007 PMID:28619474

26. Walter C, Frieden E. THE PREVALENCE AND SIGNIFICANCE OF THE PRODUCT INHIBITION OF ENZYMES. Adv Enzymol Relat Subj Biochem. Adv Enzymol Relat Subj Biochem; 1963;25: 167–274. doi:10.1002/9780470122709.CH4 PMID:14149677

27. Fachi JL, De J, Felipe S, Passariello L, Dos A, Farias S, et al. Butyrate Protects Mice from *Clostridium difficile*-Induced Colitis through an HIF-1-Dependent Mechanism. Cell Rep. 2019;27: 750–761. doi:10.1016/j.celrep.2019.03.054

28. Girinathan BP, DiBenedetto N, Worley JN, Peltier J, Arrieta-Ortiz ML, Immanuel SRC, et al. In vivo commensal control of *Clostridioides difficile* virulence. Cell Host Microbe; 2021;29: 1693–1708.e7. doi:10.1016/J.CHOM.2021.09.007 PMID:34637781

29. Martin-Verstraete I, Peltier J, Dupuy B. The Regulatory Networks That Control *Clostridium difficile* Toxin Synthesis. Toxins (Basel). Toxins (Basel); 2016;8. doi:10.3390/TOXINS8050153 PMID:27187475

30. Fuchs M, Lamm-Schmidt V, Lenče T, Sulzer J, Bublitz A, Wackenreuter J, et al. A network of small RNAs regulates sporulation initiation in *Clostridioides difficile*. EMBO J. John Wiley & Sons, Ltd; 2023; e112858. doi:10.15252/EMBJ.2022112858

31. Burns DA, Minton NP. Sporulation studies in *Clostridium difficile*. J Microbiol Methods. Elsevier; 2011;87: 133–138. doi:10.1016/J.MIMET.2011.07.017 PMID:21864584

32. Edwards AN, Mcbride SM. Initiation of sporulation in *Clostridium difficile*: a twist on the classic model. FEMS Microbiol Lett. Oxford Academic; 2014;358: 110–118. doi:10.1111/1574-6968.12499 PMID:24910370

33. Fimlaid KA, Jensen O, Donnelly ML, Siegrist MS, Shen A. Regulation of *Clostridium difficile* Spore Formation by the SpoIIQ and SpoIIIA Proteins. PLOS Genet. Public Library of Science; 2015;11: e1005562. doi:10.1371/JOURNAL.PGEN.1005562 PMID:26465937

34. Kumar N, Browne HP, Viciani E, Forster SC, Clare S, Harcourt K, et al. Adaptation of host transmission cycle during *Clostridium difficile* speciation. Nat Genet 2019; 2019;51: 1315–1320. doi:10.1038/s41588-019-0478-8 PMID:31406348

35. Donskey CJ, Chowdhry TK, Hecker MT, Hoyen CK, Hanrahan JA, Hujer AM, et al. Effect of antibiotic therapy on the density of vancomycin-resistant enterococci in the stool of colonized patients. N Engl J Med; 2000;343: 1925–1932. doi:10.1056/NEJM200012283432604 PMID:11136263

36. Lawley TD, Bouley DM, Hoy YE, Gerke C, Relman DA, Monack DM. Host transmission of *Salmonella enterica* serovar Typhimurium is controlled by virulence factors and indigenous intestinal microbiota. Infect Immun; 2008;76: 403–416. doi:10.1128/IAI.01189-07 PMID:17967858

37. Ng KM, Aranda-Díaz A, Tropini C, Frankel MR, Van Treuren W, O’Laughlin CT, et al. Recovery of the Gut Microbiota after Antibiotics Depends on Host Diet, Community Context, and Environmental Reservoirs. Cell Host Microbe; 2019;26: 650–665.e4. doi:10.1016/j.chom.2019.10.011 PMID:31726029

38. Rivera-Chávez F, Bäumler AJ. The Pyromaniac Inside You: *Salmonella* Metabolism in the Host Gut. Annu Rev Microbiol; 2015;69: 31–48. doi:10.1146/ANNUREV-MICRO-091014-104108 PMID:26002180

39. Park SY, Rao C, Coyte KZ, Kuziel GA, Zhang Y, Huang W, et al. Strain-level fitness in the gut microbiome is an emergent property of glycans and a single metabolite. Cell; 2022;185: 513–529.e21. doi:10.1016/J.CELL.2022.01.002 PMID:35120663

40. Baker SA, Rutter J. Metabolites as signalling molecules. Nat Rev Mol Cell Biol; 2023; doi:10.1038/S41580-022-00572-W PMID:36635456

41. Joseph T. Acetone/EtOH Bacterial RNA Isolation. In: labspaces.net [Internet]. 2010. Available: http://www.labspaces.net/protocols/616741283537722_protocol.pdf

42. Bender KO, Garland M, Ferreyra JA, Hryckowian AJ, Child MA, Puri AW, et al. A small-molecule antivirulence agent for treating *Clostridium difficile* infection. Sci Transl Med; 2015;7. doi:10.1126/SCITRANSLMED.AAC9103 PMID:26400909

43. Hromada S, Venturelli OS. Gut microbiota interspecies interactions shape the response of *Clostridioides difficile* to clinically relevant antibiotics. PLoS Biol; 2023;21: e3002100. doi:10.1371/JOURNAL.PBIO.3002100 PMID:37167201

44. Pensinger DA, Gutierrez K V., Smith HB, Vincent WJB, Stevenson DS, Black KA, et al. *Listeria monocytogenes* GlmR Is an Accessory Uridyltransferase Essential for Cytosolic Survival and Virulence. Kline KA, editor. mBio; 2023;14. doi:10.1128/MBIO.00073-23 PMID:36939339

45. Clasquin MF, Melamud E, Rabinowitz JD. LC-MS data processing with MAVEN: a metabolomic analysis and visualization engine. Curr Protoc Bioinformatics 2012;Chapter 14: Unit14.11. doi:10.1002/0471250953.bi1411s37 PMID:22389014

46. Melamud E, Vastag L, Rabinowitz JD. Metabolomic Analysis and Visualization Engine for LC−MS Data. Anal Chem. 2010;82: 9818–9826. doi:10.1021/ac1021166 PMID:21049934

47. Latendresse M, Karp PD. Web-based metabolic network visualization with a zooming user interface. BMC Bioinformatics; 2011;12: 176. doi:10.1186/1471-2105-12-176 PMID:21595965

